# Derivation of a minimal functional XIST by combining human and mouse interaction domains

**DOI:** 10.1101/2022.09.02.506429

**Authors:** Maria Jose Navarro-Cobos, Suria Itzel Morales-Guzman, Sarah E.L. Baldry, Carolyn J. Brown

## Abstract

XIST is a 17-19 kb long non-coding RNA critical for X-chromosome inactivation. Tandem repeats within the RNA serve as functional domains involved in the cis-limited recruitment of heterochromatic changes and silencing. To explore the sufficiency of these domains while generating a functional miniXIST for targeted silencing approaches, we tested inducible constructs integrated into 8p in a male cell line. Previous results suggested silencing could be accomplished with a transgene comprised of the repeat A which is highly conserved and critical for silencing; the repeat F which overlaps regulatory elements and the repeat E region which contributes to XIST localization by binding proteins such as CIZ1. As PRC1 is recruited through HNRNPK binding of repeats B - C - D we included a second ‘miniXIST’ comprising AFE with the mouse PID, a 600-nucleotide region of repeat B and C. Silencing of nearby genes was possible with and without PID, however, silencing more distally required the addition of PID. The recruitment of heterochromatic marks, evaluated by IF combined with RNA FISH, revealed that the AFE domains were sufficient only for the recruitment of CIZ1. However, miniXIST transgene recruited all marks, albeit not to full XIST levels. The ability of the PID domain to facilitate silencing and heterochromatic marks recruitment was unexpected, and inhibition of PRC1 suggested that many of these are PRC1-independent. These results suggest that the addition of this small region allowed the partial recruitment of all the changes induced by a full XIST, demonstrating the feasibility of finding a minimal functional XIST.

## Introduction

X-chromosome inactivation (XCI), proposed by Mary Lyon in 1961, is an essential process in eutherian mammals, where one of the X chromosomes in females (XX) is silenced to compensate for X-linked gene expression with the single X chromosome (X) of males (XY). The future inactive X (Xi) transcribes a long non-coding RNA (lncRNA), the X-inactive specific transcript (XIST) ^1^, which coats the Xi, and has a critical role in triggering the subsequent molecular events that maintain this silencing during the next cellular divisions (reviewed in ^2–6^). XCI has been widely studied in mice, where it occurs in two waves, an imprinted paternal XCI, and then later in the inner cell mass, following reactivation, random silencing takes place ^2,7^.

XCI in humans is less understood, but is known to be random in all tissues, including extra-embryonic ^8^. Human embryonic stem cells (hESC) already have an Xi, therefore human XCI initiates in early implantation, which makes its study difficult. Pluripotent stem cells (PSC) can be induced to represent the early naive stages of development and are becoming established as a potential cellular model for XCI ^7^. Studies of embryos from *in vitro* fertilization have revealed that *XIST* is upregulated from both X chromosomes, with biallelic expression of X-linked genes that are normally silenced in somatic cells ^9,10^. Similar early expression from both X chromosomes is seen in other primates ^11^.

XIST is approximately 17-19 kb long and has a modular structure, in which tandem repetitive regions labelled as A to F can be identified. Conservation of *XIST/Xist* between human and mouse varies across the repeats. XIST acts as a scaffold for the recruitment of needed proteins for XCI, and its accumulation leads to the loss of histone modifications related to the euchromatic state, such as Histone H3 lysine 27 monoacetylation (H3K27ac), and accumulation of heterochromatic marks such as Histone H3 lysine 27 trimethylation (H3K27me3), Histone H4 lysine 2- monomethylation (H4K20me1), and H2A lysine 119 monoubiquitination (UbH2A). At the present, over 80 proteins have been described to bind Xist ^6,12–14^. Distinct domains of XIST/Xist have been shown to be important for the recruitment of many of these Xi-enriched proteins and marks ^3,6^.

The repeat A in Xist comprises 7.5 copies of a 26 nucleotide core sequence at the 5’ end of Xist, with the potential to form hairpin-like structures and loops ^5,6^. This domain is necessary for silencing and interacts with SPEN ^12,15,16^, which is indispensable for the initiation of XCI, and important in recruitment of the SMRT-HDAC3 complex, that represses transcription by exclusion of RNA polymerase II ^16,17^. The A repeat region is also essential for silencing in humans ^18^ and is one of 5 hubs of protein binding identified by immunoprecipitation ^19^, binding at least 10 different (although often related) proteins. The F region was the second such hub, binding 10 proteins including the HNRNPK and U proteins, which are enriched in aggregates across the BCD region. The B repeats are split in humans, and the C repeat region of mouse is less than a single copy in humans, while the D repeat region is expanded in humans to include 14 repeats, as well as additional degenerate repeats in the region between C and D ^20^. HNRNPK is a DNA/RNA binding protein, and it interacts with the mouse repeat B, specifically with the Polycomb Interaction Domain (PID), a 600 nucleotide region, that is essential for the recruitment of Polycomb-repressive complexes PRC1 and PRC2 ^21–23^. In humans, HNRNPK binds the B, C and D domains, and the protein is additionally implicated in cancer or rare disease ^14,22,24^. PRC1 catalyzes UbH2A, whereas PRC2 catalyzes H3K27me2/3 and they are vital in the maintenance of XCI ^6,25,26^. These complexes are built up of different subunits, and the traditional model where Xist directly recruits PRC2 is being questioned, with an alternative proposal arguing that PRC2 binding is important for the improvement of its own functions. PRC1 is recruited by HNRNPK, which acts as an adaptor protein ^26^, moreover HNRNPK depletion decreases UbH2A deposition ^25^.

CIZ1 is a nuclear matrix protein, and its binding to repeat E contributes to Xist localization ^12,21,27^. Different studies have suggested that repeat domains C, D, and E appear to be important for the localization of Xist ^27–30^. HNRNPU is reported to contribute to XIST localization in at least some cells ^31,32^. Interestingly, pathogenic variants in CIZ1 are the basis for the etiology of adult-onset primary cervical dystonia ^33^. The E repeat region is another protein binding hub, binding a number of co-operatively interacting proteins that contribute to phase separation ^34,35^.

Unlike autosomes and the active X (Xa), which are organized into A/B compartments and topologically associated domains (TADs), the Xi is organized in 2 megadomains, separated by a macrosatellite repeat, *Dxz4*. SMCHD1 is a non-canonical structural maintenance of chromosome (SMC) protein that is required for the maintenance of XCI. Originally identified during a screen for chromatin remodelers, murine Smchd1 shows female-specific lethality ^36^. SMCHD1 recruitment is dependent on the UbH2A mediated by PRC1 ^37^, and in male *Smchd1*−/− embryonic stem cells there is upregulation of CpG hypermethylation of the *Pcdh* genes that are important for neuronal diversity, and also aberrant expression of *Hox* genes. Knockdown of human *SMCHD1* leads to Xi decompaction and erosion of heterochromatic silencing ^38,39^. Further, mutations in *SMCHD1* are a cause of facioscapulohumeral muscular dystrophy type 2 (FSHD2) and arhinia; however, surprisingly, in female patients, XCI is preserved. Nevertheless, in a subset of patients with arhinia, upregulation of the *PCDHA* cluster is reported, similar to the mouse knockouts ^40,41^.

Recent studies from our lab using an inducible human XIST transgene in a male somatic cell line to study the initiation of heterochromatic silencing by XIST have shown differences between polycomb recruitment in humans from mice, and that each region tested, repetitive or not, was critical for recruitment of at least one feature of XCI. These deletion studies demonstrated that the A and F regions were critical for silencing as well as H3K27me3 and ubH2A recruitment. The E repeat was the region required for most features, and together the end of the XIST inducible construct (including E) was required for each feature studied in our previous analysis. XIST has been tested as a potential therapeutic agent, by silencing the extra chromosome 21 in Down syndrome ^42^. Therefore, a minimal functional XIST, that is sufficient to induce silencing and heterochromatin formation, could contribute to chromosome therapy. To identify which domains of XIST are necessary and sufficient to provide functionality to XIST, we assembled two minimal XIST transgenes, the first is 4.5 kb and includes the A, F and E repeats, and a second one that is 5.1 kb, with the A, F and E repeats plus the PID region from mouse.

## Results

### Mouse PID partially rescues lack of silencing by human XIST transgene lacking B - D repeat region

Induction of an XIST cDNA integrated into an FRT site on chromosome 8p results in both silencing of flanking 8p genes and recruitment of heterochromatin marks^43^. Assessment of deletions spanning this full XIST reinforced that the 5’ end of XIST including the conserved A repeats was critical for silencing ^44^. While the 5’ A repeats are essential for silencing, they are only sufficient to induce local silencing ^45^. The region including the E repeats binds matrix proteins and thus may allow the spreading of A-repeat-mediated silencing. We hypothesized that a construct containing the 5’ 1.9 kb, which includes the A repeats and the regulatory elements surrounding the F/P2 region ^46^, as well as the 3’ end of the XIST cDNA from the end of exon 1 would be able to silence flanking genes. Therefore, we generated a construct containing these regions, named AFE. Since the B-C-D regions are involved in polycomb recruitment in humans, whereas in mice the PID region of B-C is required ^25^, we also generated a miniXIST containing AFE with the mouse PID between the 3’ and 5’ regions to test whether PID would be sufficient for polycomb protein recruitment in humans. For our analyses, we included the previously studied full XIST transgenes as a control (Figure 1A).

**Figure 1:**
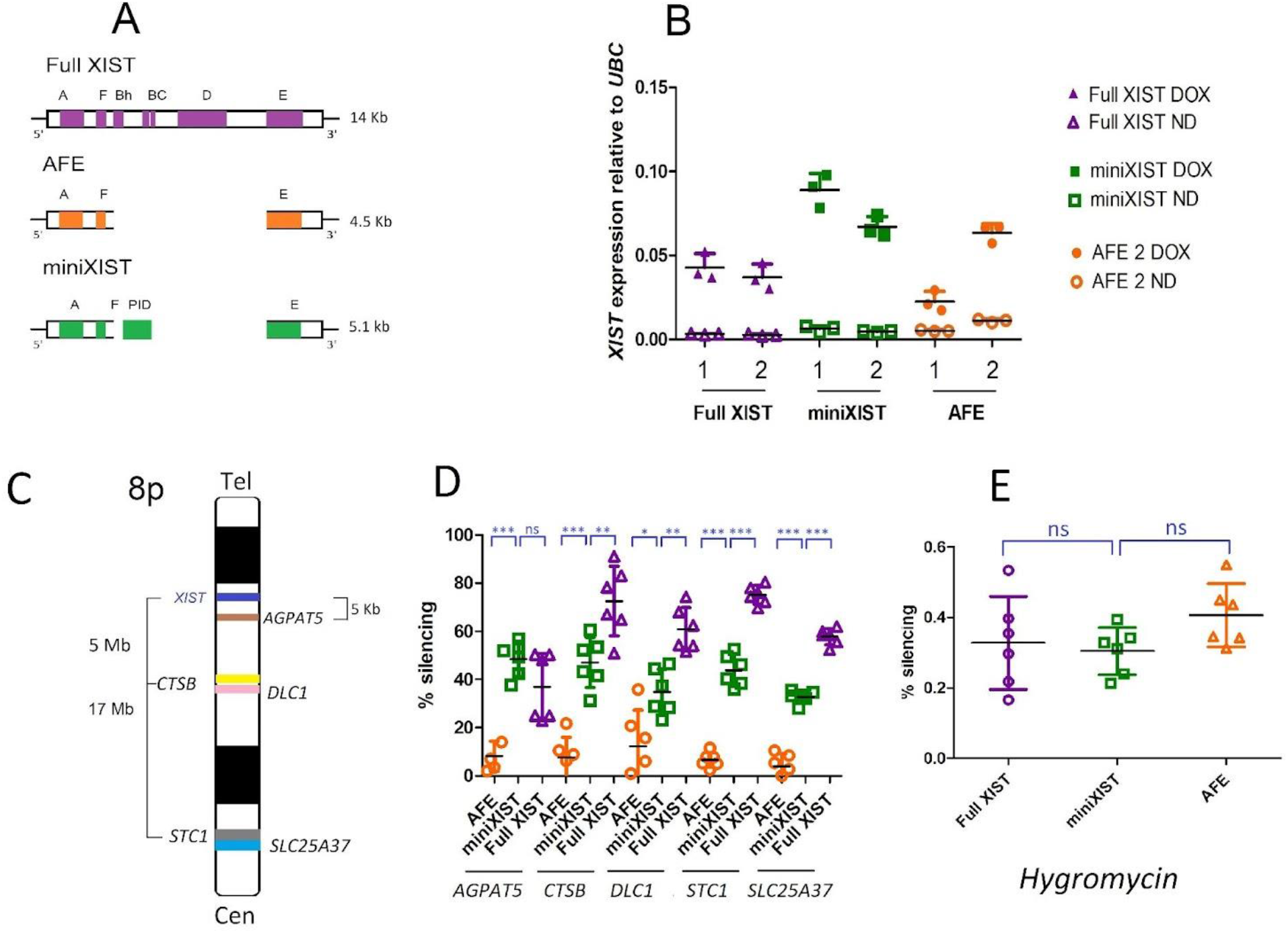
The ability of minimal XIST constructs to induce silencing. A. Schematic of the AFE and miniXIST constructs relative to Full XIST that has been previously studied. B: qPCR showing XIST expression for two clones of each construct along with full XIST relative to the internal control *UBC*. Data is average of three replicates. C and D: Silencing induced by the three constructs for five genes, *AGPAT5* at 10kb, *CTSB* and *DLC1*, which are 5 MB and *STC1* and *SLC25A37* at 17 MB away from the integration site. E: silencing of *Hygromycin* gene, which is at 0.6 kb, by qPCR relative to the *UBC* internal control.

To enable comparison of single-copy integrations at the same genomic site, all constructs were analyzed following FRT-mediated integration into the 8p of the HT1080 male fibrosarcoma cell line. After 5 days of induction with Doxycycline (DOX), which results in the removal of the TetR repressor from the CMV promoter present in the constructs, we evaluated two transgenic lines per construct, with the experiments performed in triplicate. We confirmed *XIST* expression in all the transgenic lines by qRT-PCR, noting some variability in the level of XIST induction between different lines (Figure 1B). To examine the spread of silencing we selected five genes with expressed SNPs at varying distances from the FRT integration site for the transgene: *AGPAT5* (10 kb), *CTSB* (5 Mb); *DLC1* (6Mb); *STC1;* and *SLC25A37* (17 Mb). These genes had previously been shown to be partially silenced by XIST induction, and the percentage of allelic silencing was calculated by measuring the relative expression of SNPs in DOX and NoDOX samples by pyrosequencing (Figure 1C).

The AFE construct exhibited low allelic silencing with 8.1%, 7.6%, 12.3%, 6.6% and 3.9% silencing for *AGPAT5*, *CTSB, DLC1, STC1* and *SLC25A37* respectively. In contrast, the full XIST transgene demonstrated 38%, 72.5%, 60.8%, 75.1% and 57.7% for *AGPAT5*, *CTSB, DLC1, STC1* and *SLC25A37*. We further tested whether the addition of the PID to AFE would improve silencing. The miniXIST transgenes showed 48.5%, 47%, 34.7%, 43.8%, 32.7% silencing (Figure 1D). Unexpectedly, it appears that AFE only induced very weak silencing, yet miniXIST consistently had an intermediate level of silencing for *CTSB, DLC1, STC1* and *SLC25A37*, but never achieved the high level of silencing of full XIST. The FRT integration site on 8p is within the *AGPAT5* gene and the percentages of silencing for miniXIST and Full XIST were similar. The *Hygromycin* gene is present in the constructs, therefore we tested its silencing by qPCR. Interestingly, the percentage of silencing induced by all constructs for this nearby gene was not significantly different from each other (Figure 1E).

### Addition of PID region also induces partial chromatin structure changes

In addition to examining silencing, we wanted to assess whether the AFE construct would recruit chromatin changes and, further, if there was a difference in this recruitment between AFE and miniXIST that could explain the substantial improvement in the ability to silence endogenous genes. The cells were grown on coverslips for Immunofluorescence-Fluorescence in situ hybridization (IF-FISH) with 5 days of DOX induction. In addition to the six transgenic lines with the three constructs, we included a female cell line, IMR-90, as an additional positive control. We performed IF assays for the heterochromatic marks and related proteins H3K27me3, H4K20me1, UbH2A, MACROH2A, SMCHD1, CIZ1, and the euchromatic mark H3K27ac. The IF was combined with RNA FISH for XIST. After visual confirmation of both signals, we used a quantitative approach to evaluate the presence of the marks or proteins (in red) in the same region where XIST is visualized (in green). The calculation of a Z score in individual cells allowed us to quantitate the level of enrichment for the red signal where the green one (XIST) had its highest intensity (Figures 2A and supplementary 2-8). A Z score value of 1 means that the red signal is 1 standard deviation higher where XIST is maximal. The blinded analysis of 200 cells per construct revealed that for the same mark some cells were enriched at the XIST signal and others not. H3K27me3 and UbH2A are catalyzed by PRC2 and PRC1 respectively, and previous deletion studies had suggested that these complexes are recruited by different human XIST domains ^18^. PRC1 recruitment to Full XIST required the BC region as well as regions including the A, D and 3’ domains; whereas in mice this complex only requires the PID domain. PRC2 recruitment required the human F-Bh and E domains; whereas, in mice, PRC2 recruitment has been reported to be downstream of PRC1 recruitment ^25^. Thus, we anticipated that the AFE construct might recruit at least H3K27me3, meanwhile, the miniXIST constructs would be able to recruit both marks. Intriguingly, more than half of the AFE construct cells had a negative Z score, showing overall averages of 0.02 for H3K27me3 and 0.10 for UbH2A. When we analyzed the cells with the miniXIST construct we again found cells with both positive and negative Z scores. When the miniXIST averages were calculated they were lower than either full XIST or IMR-90. The averages for H3K27me3 were 0.46 versus 1.17 found in full XIST (p<0.0001). UbH2A averages were 0.39 versus 0.99 for full XIST (p<0.0001). The second positive control, IMR-90, displayed averages of 1.79 and 1.15 for both marks respectively (Figures 2B and 2C). Nevertheless, both positive controls also had some negative Z score cells. These averages suggest that the AFE construct is not able to recruit PRC2. Meanwhile, the miniXIST construct improves the Z score averages not only for PRC1 but also for PRC2, however, the averages were still lower when compared with the positive controls.

**Figure 2:**
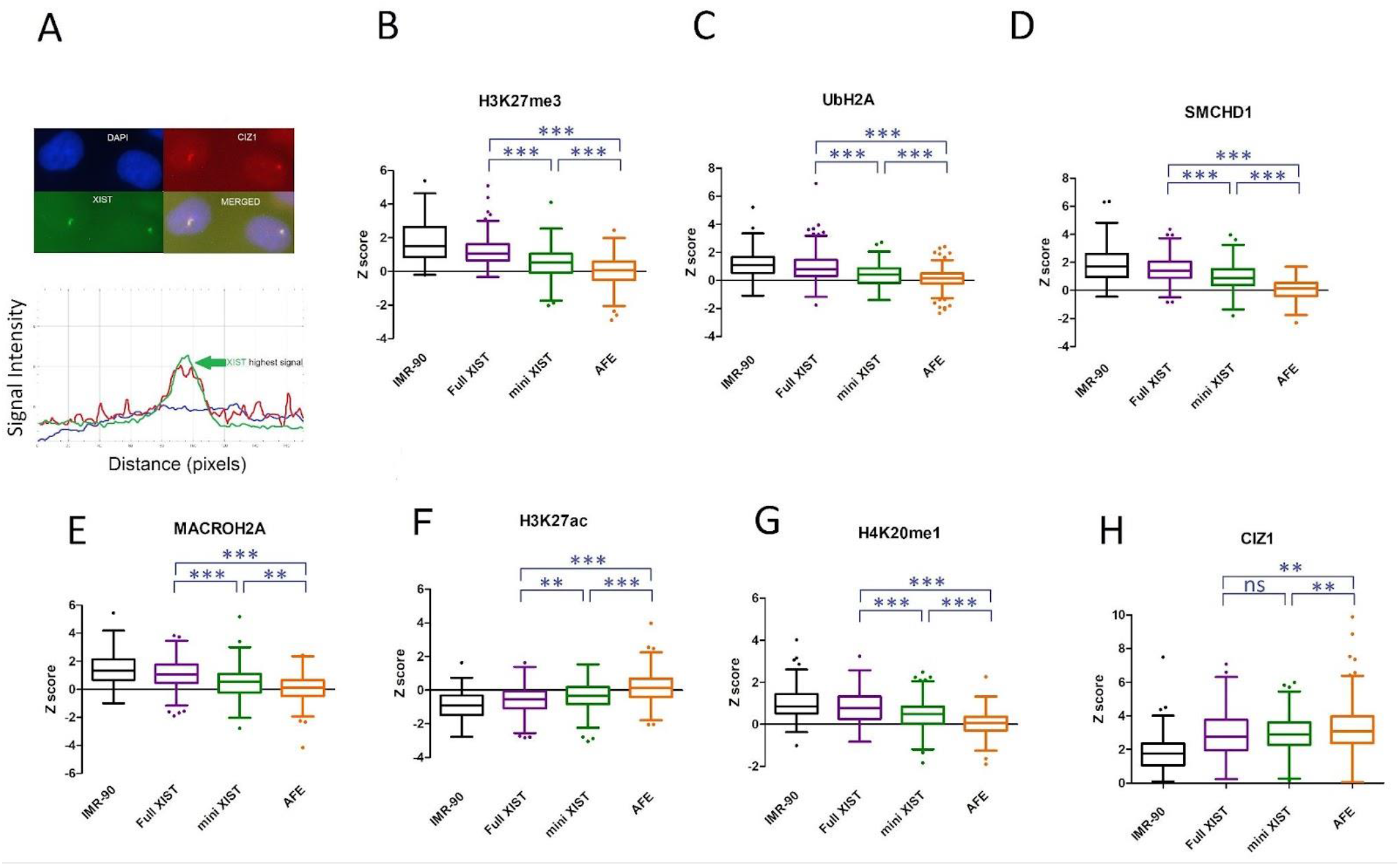
The miniXIST construct promotes partial recruitment of heterochromatin and loss of a euchromatic mark. A: Example of the images obtained from RNA FISH combined with IF for protein CIZ1, DAPI (blue), XIST (green), CIZ1 (red) and the merged image for the three channels. The graphic illustrates that the XIST peak in green coincides with the peak for CIZ1 in red, showing the co-localization of both signals. B, C, D, E, F, G, H: Z score calculated for three heterochromatic marks UbH2A, H3K27me3, H4K20me1, loss of one euchromatic mark H3K27ac, and three related proteins CIZ1, macroH2A, and SMCHD1, for the three constructs (2 clones each) and IMR-90 cell line, 100 cells/construct/mark were analyzed, t-test was calculated, the asterisks represent the significance, corrected P-value <0.0001***, 0.0001-0.01**, 0.01-0.05*, >0.05 ns.

Recent work from the laboratory demonstrated that PRC1 is also needed for SMCHD1 recruitment and PRC2 for MACROH2A recruitment by XIST ^18^. Thus, we next evaluated both proteins. Similar to previous results, the AFE construct Z score averages were 0.07 and 0.09 for SMCHD1 and macroH2A, suggesting that this construct was not recruiting either of them. The miniXIST construct averages showed better recruitment, with averages of 0.95 and 0.44 for SMCHD1 and macroH2A. Full XIST and the IMR-90 cells displayed the highest averages. For full XIST, the average for SMCHD1 was 1.48 and 1.11 for macroH2A. IMR-90 had averages of 1.93 and 1.44 respectively (Figures 2D and 2E). Although both the AFE and the miniXIST constructs had different Z score averages, we reiteratively found cells with negative and positive Z scores.

We also examined the loss of the euchromatic mark H3K27ac. This mark had an inverse trend in the constructs. The AFE construct retained the highest average (0.13), suggesting no depletion at the XIST locus. The miniXIST construct revealed a lower average of −0.34. Full XIST and IMR-90 averages were the lowest, although some cells showed a positive Z score, with averages of −0.59 and −0.93 respectively (Figure 2F). Even though the miniXIST construct had some cells with a positive Z score, the average was negative. H4K20me1 is another heterochromatic mark found on the Xi, which is deposited by SETD8 ^47^. The Z-score averages found were 0.05 for the AFE construct and 0.46 for miniXIST. Again, the controls full XIST and IMR-90 averages were the best showing 0.82 and 1.04 respectively (Figure 2G). Unexpectedly, this mark also exhibited improvement when the PID region is present. Finally, we examined CIZ1. This protein binds the repeat E ^48^, which is present in all constructs. Contrary to the rest of the studied marks, CIZ1 was recruited for all constructs with averages of 3.27, 2.92, 2.96, and 1.82 for AFE, miniXIST, full XIST, and IMR-90 respectively (Figure 2H), showing no improvement with the presence of additional domains of XIST. Overall, these results suggest that the AFE construct fails to establish XCI-related chromatin changes with the exception of CIZ1, whereas miniXIST had an improvement for all the analyzed marks, demonstrating an intermediate level of recruitment when compared to full XIST and the IMR-90 cell line.

### PRC1 inhibition did not affect PRC2 related marks or gene silencing

The IF RNA FISH results were highly consistent for all marks and proteins, and unexpectedly, suggested that the PID region had a positive impact on the recruitment of both PRC1 and PRC2, as well as related proteins. Therefore, our next question was whether this recruitment was mediated through PRC1 enzymatic activity. Thus, we evaluated the effect of PRC1 inhibition on the full and miniXIST constructs to corroborate if the other marks were dependent on UbH2A recruitment. PRT4165 is a potent PRC1 inhibitor ^49^, having a direct effect on H2A ubiquitylation. First, we tested the effect on UbH2A recruitment by using IF RNA FISH with cells grown with DOX induction and with or without PRT4165 for 5 days. We used 45 uM PRT4165 as lower concentrations had a limited impact on UbH2A at the 8p chromosome, and higher doses impacted cell survival. At this concentration, a significant reduction in Z scores for UbH2A recruitment was found for all constructs (Figure 3A). IMR-90 and full XIST presented Z score averages of 1.85 and 1.21 without the inhibitor and 0.91 and 0.73 with the inhibitor respectively. The miniXIST construct showed 0.39 and 0.07 without and with the inhibitor (Figure 3A), supporting that the inhibitor at 45uM concentration was affecting the recruitment. Because SMCHD1 requires PRC1 activity we next evaluated this protein. Both controls and miniXIST displayed a significant reduction in the Z score averages. IMR-90 cell line and full XIST went from 1.83 and 1.08 to 0.31 and 0.31 respectively, whereas the miniXIST construct exhibited a decrease from 0.56 to −0.16 (Figure 3B). Subsequently, H3K27me3 and macroH2A were evaluated. The analysis of H3K27me3 revealed that even though there was a slight decline in the Z score averages in the control lines, this was not significantly different. The same was true for the miniXIST construct which did not have a change in the recruitment of this mark (Figure 3C). Finally, we investigated the recruitment of MACROH2A. This mark appears to be PRC2-dependent ^18^. Similar to H3K27me3, all the constructs including the miniXIST showed no difference in the Z scores averages when the PRC1 inhibitor was added (Figure 3D). The results suggest that when PRC1 is inhibited in the miniXIST construct only UbH2A and SMCHD1 are impacted, but not the chromatin changes induced by PRC2; despite the addition of the PID leading to partial recruitment of those marks.

**Figure 3:**
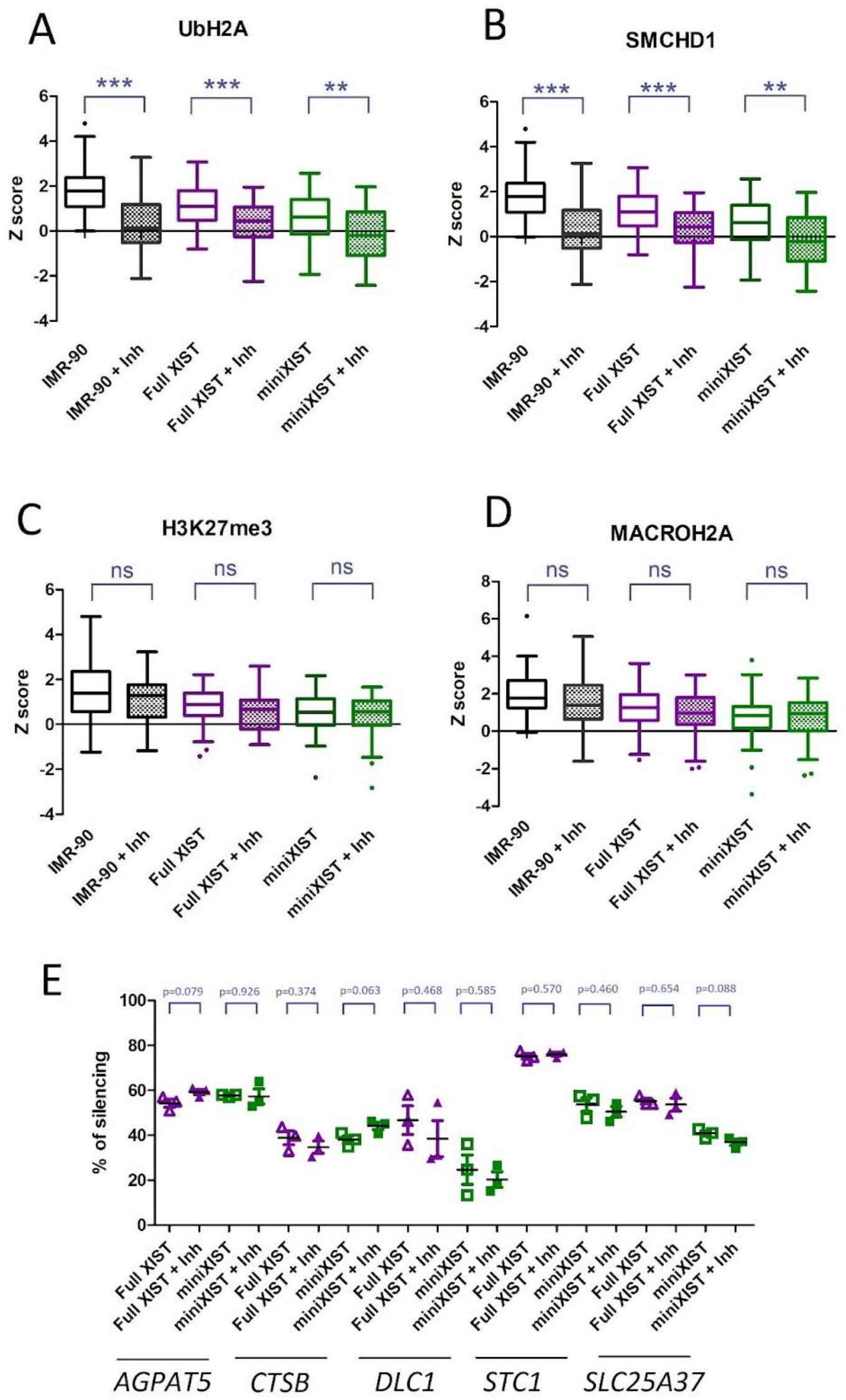
PRC1 inhibition had an impact on UbH2A and SMCHD1 but not on PRC2 recruitment. A and B: UbH2A and SMCHD1 Z scores declined when the PRC1 inhibitor was applied. C and D: for H3K27me3 and macroH2A the PRC1 inhibitor did not show an effect. E: Silencing of *CTSB, DLC1, STC1,* and *SLC25A37* was not affected by the addition of PRC1 inhibitor PRT4165 when compared with the samples only induced with doxycycline

Finally, to examine whether there was any impact on silencing, we used pyrosequencing of cDNA from the *AGPAT5, CTSB*, *DLC1*, *STC1,* and *SLC25A37* genes. No significant difference in silencing was detected in the analysis of triplicate samples. Full XIST construct silencing averages were 53.9% and 50.7% and the miniXIST had 39.2% and 37.7% for cells without and with the inhibitor at 45 uM concentration respectively (Figure 3E).

### HNRNPK binds the miniXIST construct upon addition of the PID region

In mice, the PID region recruits HNRNPK, which is then proposed to recruit PRC1. HNRNPK binds C-rich stretches ^23,25,50^, and the B-repeat part of PID is very CCC rich, although the part derived from the C-repeat also contributes potential binding sites. In humans, only a partial single copy of the region paralogous to the mouse C repeat is present and has no CCC sequence. Furthermore, B and Bh combined would not give as many CCC as PID, thus in humans, D may also be contributing to HNRNPK recruitment (Figure 4A). To verify that the PID region was recruiting HNRNPK, as is observed in mice, we performed UV-crosslinked RNA immunoprecipitation (RIP) for full XIST, miniXIST, and AFE. The cells were induced for 5 days with DOX, and then UV-crosslinked prior to precipitation using antibodies for HNRNPK and IgG as a control. Subsequently, RNA extraction of the lysate and qPCR revealed the presence of XIST in the pull-down. The AFE construct was unable to pull-down XIST, while the average pull-down percentages were 7.3% for full XIST, 4.2% for miniXIST and 0.8% for AFE (Figure 4B). We found substantial variability in the pull-down percentages, such that a T-test comparison of Full XIST and miniXIST revealed that they were not significantly different from each other. This finding supports that the PID region is recruiting HNRNPK in human cells.

**Figure 4:**
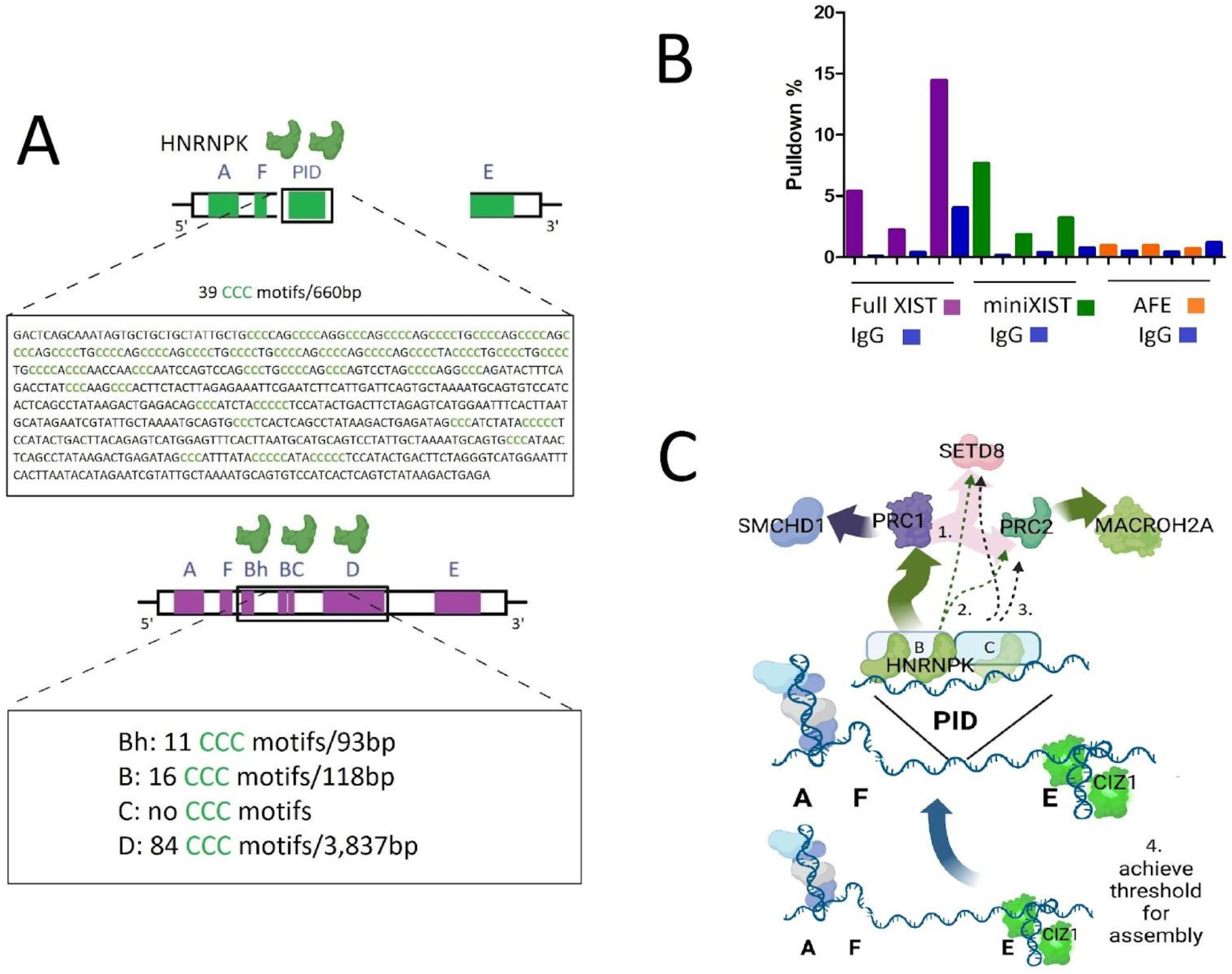
PID region recruits HNRNPK. A: PID mouse region (660 bp) is rich in CCC motifs, which could be important for HNRNPK binding. The possible candidate regions for the recruitment of HNRNPK in humans are the Bh, B, C and D domains. Of those, Bh (93 bp) and B (118 bp) are small in size and encompass several CCC motifs. B: pulldown percentages revealed that HNRNPK is binding XIST in both full XIST and miniXIST, whereas AFE was not able to recruit this protein. C: Possible ways that the PID region could be using to recruit the different heterochromatic marks. 1. The part of PID region that encompass domain B in bound by HNRNPK which in turns recruits PRC1 and SMCHD1. 2. HNRNPK also recruits PRC2 and MACROH2A. 3. The domain C present in the PID region recruits direct or trhough unidentified proteins PRC2 and MACROH2A. 4. The addition of the PID region is crossing a necessary threshold to the recruitment of all heterochromatic marks.

## Discussion

The functional domains of XIST are not yet fully understood; however, studies have identified many interacting proteins (reviewed in ^51^) with binding in humans identified by RNA immunoprecipitation aggregating in 5 hubs ^19^. Because of the capacity of XIST to trigger sufficient chromatin changes to silence an entire chromosome, it has potential for chromosome therapy ^42^. However, the size of a full version of XIST is unwieldy. Thus, based on the current knowledge about the different functions of each XIST domain, we tried to generate a smaller construct that could recapitulate all the XIST functions. Domain A is crucial for silencing by recruiting proteins at the initiation of XCI in both, mice and humans ^5,6,15,44^, and domain E is relevant for XIST/Xist localization also through protein interactions ^12,28,29,48^. While A and E domains are clearly critical, our recent results suggested that the functional pathways utilized in human somatic cells may differ from those described in mouse differentiation (reviewed in ^7^). These prior studies used deletions to identify necessary regions, but we wanted to test if such regions were sufficient to induce silencing and/or chromatin changes. Therefore in this work, we tested two new XIST transgenes: a 4.5 kb construct with the AFE domains; and a second one, 5.1 kb long with AFE plus the mouse PID region. While our recent study suggested that AFE might be sufficient for silencing, the mouse PID was recently recognized as sufficient for PRC1 recruitment ^25^. We induced expression of both constructs and the previously studied full XIST from a single integration site at 8p location. This allowed us to analyze if some principal XIST-induced events of XCI were differentially triggered by the constructs.

We started by testing silencing, which was anticipated to happen for both transgenes. It is known that repeat A is necessary for silencing and sufficient for local silencing ^45^, and we did observe silencing of the Hyg gene by all constructs. Surprisingly, the AFE construct was not able to spread substantial silencing to any of the other genes, no matter the distance between the insertion site and the tested gene. *AGPAT5* was silenced by both the miniXIST and the full XIST to similar extents. However, for the other four flanking genes, we observed a clear trend with the control full XIST having the greatest silencing. Thus, we observe that AFE can silence only a very short distance; while miniXIST is equivalent to Full XIST only for the encompassing gene for which the promoter is only 10 kb away. Despite the ability of miniXIST to silence these nearby genes as well as full XIST, the silencing of multiple genes 5 to 17 Mb away was consistently reduced by approximately half.

In a previous study in the laboratory, it was found that the 3’ end of XIST was necessary for silencing, and also for UbH2A and SMCHD1 recruitment ^18^. Based on previous constructs ^42^, and the fact that exon 7 is alternatively included, we reduced the 3’ of our constructs to include only exon 6 up to the internal splice site. As miniXIST was able to silence and recruit UbH2A and SMCHD1, albeit only partially, we believe the essential 3’ region is contained in the 100 bp after the E repeat domain, which is included in miniXIST (as well as AFE). However, even incorporating this domain, silencing was not equivalent to full XIST, suggesting that some aspect of heterochromatin remodelling had not been achieved.

Since PID is known to recruit PRC1, we looked at chromatin changes promoted by miniXIST. The findings were strikingly similar to silencing. Although we observed some variability in the Z score averages for different clones, the recruitment patterns within the two clones of the same construct were consistent with each other. PRC1 recruitment by miniXIST was only 42% of that seen for full XIST. We had anticipated that the PID region would recruit UbH2A, as it does in mice. In mouse, recruitment of SMCHD1 is dependent on the UbH2A mediated by PRC1^37^. In humans, SMCHD1 is also dependent on UbH2A presence ^18^, and we also confirmed partial recruitment for this protein. Among the domains needed for SMCHD1 recruitment, besides B and C domains, are repeat D and the region 3’ to repeat D that includes exons 1 to 6 ^18^. The miniXIST sequence does not carry that large region; however, it encompasses the internal exons 2 to 5 and exon 6, suggesting that the minimal required region to recruit SMCHD1 could be smaller than thought. H3K27me3 seems to require the presence of F, Bh and E domains in humans. When PRC2 recruitment was tested, we also found an intermediate level compared to full XIST. The current model in mouse suggests that PRC2 is recruited by PRC1, therefore we thought this mark could be dependent on PRC1. Nonetheless, when we performed the inhibition of PRC1 we did not see any changes in PRC2 recruitment. Thus, it is likely that H3K27me3 partial enrichment by PID is not being recruited via PRC1. As previously mentioned, HNRNPK binds the CCC motifs present in the PID region. Those motifs are present mainly in the first half of PID (Figure 4A). It is possible that low complexity motifs in the second half or that secondary structures formation promoted by this part of PID could be recruiting PRC2 directly or through some unknown protein (Figure 4C). In humans, macroH2A recruitment depends on the F, Bh, D and E repeats ^18^. The addition of PID also improved the recruitment of macroH2A, nevertheless, this recruitment was also partial. Our findings suggest that the PID region is at least partially replacing the function of the D domain for the recruitment of this mark. Surprisingly, H4K20me1, which is a mark unrelated to PRC1 or PRC2 was also partially recruited by the addition of the PID region. H4K20me1 depends on SETD8 during XCI in mice. Inducible constructs lacking the B and C repeats in mice do show an absence of H4K20me1^52^. It is not known if the recruitment of H4K20me1 is SETD8-dependent or if other proteins are involved in its enrichment in this XCI context. CIZ1 is dispensable for XCI establishment but crucial for maintenance by retaining H3K27me3 and UbH2A onto the Xi ^35^. The repeat E is needed and sufficient for CIZ1 recruitment, and this was confirmed by our results. All constructs, including AFE, were able to promote the enrichment of CIZ1. Additionally, we tested the loss of H3K27Ac, a euchromatic mark. While all constructs lost H3K27Ac, the AFE construct had the highest Z score averages when compared with miniXIST and controls. This result is consistent with the previous findings about the AFE construct, which is not sufficient to trigger the expected XCI events.

Even though the addition of PID made a difference in both silencing and chromatin changes, this region was not as efficient as full XIST. This was surprising in two regards - both that one region was able to impact so many features, and also that no feature was fully recruited.

A striking aspect of our findings was that multiple features (silencing, H3K27me3, ubH2A, H4K20me1, MACROH2A and SMCHD1) were all recruited by miniXIST to approximately 50% of the levels seen with the full XIST construct. We consider several alternatives for such a large impact from a small domain. First, it seemed possible that similar to mouse, the PID recruited HNRNPK which recruited PRC1 to lay down ubH2A that enabled SMCHD1 and H3K27me3 and H4K20me1 recruitment, enabling silencing ^23,25,37,53^. However, inhibition of PRC1 did not impact H3K27me3 recruitment. Even though the interdependence between PRC1, PRC2 and *Xist* is well known, and a current popular model supports the dependence of PRC2 on PRC1 acting first, it has also been reported that despite knockout of PRC1 an important percentage of cells kept the H3K27me3 mark, suggesting PRC1-independent recruitment ^54^. Thus, a second alternative would be that the PID region, either through direct binding or through binding HNRNPK or another protein, independently enables PRC1 and PRC2 recruitment (Figure 4C). PRC2 is known for binding directly to different RNAs ^26^. PID mouse region recruits other proteins besides HNRNPK, such as SAP18, RYBP, PCGF5, RNF2/RINGB1 ^55^, thus there is support that the partial recruitment that we identified for the miniXIST construct was not dependent on PRC1. For the other proteins, previous studies in mice ^37,53^ and humans ^18^ have shown that SMCHD1 recruitment relies on PRC1, and our previous study in humans demonstrated that MACROH2A was dependent on PRC2 activity. Thus, the recruitment of MACROH2A and SMCHD1 is not surprising given that PRC2 and PRC1 were recruited. H4K20me1 recruitment has been reported to require the BC region, and occur concurrently with H3K27me3 but with different dynamics and gene distribution ^52^. There is no report of SETD8 being recruited by PRC1 or PRC2, therefore it is possible that H3K27me3 and H4K20me1 are initially triggered by the same region, or the same factor, and then different proteins are needed for placing or keeping the marks. A third option could be that the addition of the PID region could be crossing a necessary threshold for protein binding and as a consequence, it would be inducing the partial functionality that we documented (Figure 4C). It is known that Xist forms foci with proteins such as SPEN and HNRNPK, and those foci are phase-separated, and silence nearby and distant genes via chromatin compaction ^56^. Recent work has further shown that there are cell-type specific interactors with XIST ^57^. The large 17-19 kb XIST may be selected for regions that may be important in different cell types. It may be that the PID region is particularly important in maintaining XIST functionality in somatic cells, thus explaining the unexpected recruitment induced by adding the PID mouse region.

Although the PID returned binding of PRC1/PRC2/MACROH2A/SMCHD1/H4K20me1, strikingly all of those marks were only recruited to about one half of the levels to which they were recruited with full XIST, suggesting that there must be some elements missing that did not let miniXIST construct to have the same efficiency as full XIST. One possibility is that the miniXIST construct has all the necessary elements, but the amount of recruited proteins and marks is not sufficient. The PID mouse region is part of the B and C domains in mice ^25^, and is proposed to act through recruiting HNRNPK via CCC motifs. In humans, the B domain is divided into Bh and B domains, whereas the C domain is smaller than the one present in mice ^19,51^. Bh and B are diverged from each other but are rich in the CCC motif. Therefore,it may be that the Bh and B domains are the equivalent region for PID. There are fewer CCC motifs in B and Bh than in the PID region, suggesting PID could adequately replace the ability to recruit HNRNPK. However, in humans, HNRNPK was shown to bind across the B, C and D domains ^19^. While the human region paralogous to part of a mouse C repeat does not contain CCC motifs, the D domain is also rich in CCC motifs (Figure 4A), although those motifs are more dispersed in this big domain compared to B and Bh. Thus, this number or distribution of the CCC motifs may be necessary for sufficient HNRNPK recruitment to achieve the silencing seen with full XIST. A second possibility is that the domains not present in the mini XIST construct are crucial to refine and complete the recruitment at sufficient levels for all the needed marks. Deletions of Bh and B and C demonstrated that Bh is important for PRC2 recruitment, whereas B and C are needed for PRC1 recruitment ^18^, thus perhaps Bh, B and C have unique functions in human XIST. Furthermore, the D repeat region is also diverged in humans ^20^, thus it may have acquired functions other than recruitment of HNRNPK.

A smaller XIST could improve its utility for chromosome therapy, and furthermore, would lead to a better understanding of the individual functions of the different XIST domains. In this work, we generated an inducible *XIST* transgene of only 5.1 kb that was able to promote partial silencing and recruitment of heterochromatic marks. While the efficiency was not as good as expected for a full XIST version, our finding highlights the possibility of generating a minimal functional XIST.

## Materials and Methods

### Generation and culture of cell lines

Cloning of full XIST has been previously described ^58^. To generate AFE, the A-F region was amplified from full XIST with primers (see Table S1) with added restriction enzyme sites and the 1.862 bp KpnI-NotI fragment cloned into pcDNA5 FRT TO containing the 2,726 kb XhoI-ApaI fragment flanking the E repeat generated by amplifying qXIST10 (with Apa/NarI) and C7-6 (with AgeI/XhoI). For the generation of miniXIST, the 702 bp mouse PID region was synthesized by GENEWIZ and excised from the pUC57 vector with NotI and XhoI sites for integration between the AF and E regions. The two new constructs were recombined via FRT sites into the 8p FRT site on chromosome 8p of the HT1080 cell line. This site of insertion has previously been shown to allow inducible full XIST expression and silencing of nearby genes and recruitment of heterochromatic marks as well, after 5 days of induction with Dox ^43^. Two independent clones of each construct were grown in DMEM media plus 10% Fetal Bovine Serum, 5 ml of L-Glutamine, non-essential amino acids and 5 ml of Penicillin-Streptomycin. XIST expression was induced with Doxycycline (1 ug/ml of media) for 5 days.

### RNA extraction and RT-qPCR

RNA extraction of all samples was done with TRIzol (Invitrogen) reagent following its protocol, then the RNA was treated with DNAse (Roche) l U/ul per sample, for 20 minutes at 37° C, and then at 75° C to deactivate the enzyme. This RNA was converted to cDNA by reverse transcription using M-MLV Reverse Transcriptase (ThermoFisher). Each reaction consisted of 4ul of first strand buffer, 2 ul of 0.1uM DTT, 2ul of dNTPs at 1.25uM, 1ul of random hexamers, 1ul of M-MLV and 2 ug of the RNA sample. The final volume was 20 ul, using DEPC-treated H_2_O. Incubation was at 42 °C for 2 hours, followed by 95 °C for 5 minutes. cDNA was stored at −20 °C. qPCR reactions consisted of 4ul of 5x Buffer, 1.2 ul of 25 uM MgCl2, 0.16 ul of dNTPs 25 uM, 0.2 ul of primers (forward and reverse combined), 0.1ul of Go Taq G2 DNA polymerase (Promega), 1 ul of EVA Green 20x (Biotium), and 1.5 ul of the template. The qPCR was done with a QuantStudio 3 (Thermo Fisher) thermocycler, using MicroAmp Fast Optical 96-Well Reaction Plates (Applied Biosystems), and the PCR program was 40 cycles of 95 C for 30 seconds, followed by 60 °C for 30 seconds and finally 72 °C for 1 minute. The expression of the constructs was compared with the endogenous expressed gene *UBC*. The relative expression to *UBC* was calculated using the ΔΔCt method.

### Pyrosequencing

cDNA was used for PCR for the genes *AGPAT5, CTSB, DLC1, STC1* and *SLC25A37* using the following conditions 2.5 ul of 10x Buffer, 0.2 ul of dNTPs 25 uM, 0.75 ul of MgCl2 50 uM, 0.5 ul of primers forward and reverse 25nmol (Table S2), 0.125 ul of Taq Polymerase (ThermoFisher), 19.5 ul of dd H_2_O and 1 ul of the PCR product of each gene. PyroMark Q96 Systems (QIAGEN) was used. 10ul of the PCR product was mixed with 38 ul of Binding Buffer, 2 ul of Streptavidin Sepharose High Performance beads (GE Healthcare) and 35 ul of H_2_O. This mix has to be shaken at 1400 rpm for at least 5 minutes. Meanwhile the sequencing primers are prepared by adding 0.1444 ul of each primer at 25 nmol/ul with 11.856 ul of Annealing Buffer into the PyroMark optical plate. Next, the four workstation containers are filled with this order, 1.- 100 ml of 70% Ethanol, 2.- 100 ml of Denaturing Buffer, 3.- 120 ml of Wash Buffer, 4.- 100 ml of H_2_O. The workstation comb is rinsed with H_2_O, then the PCR mix is sucked up with the comb using the vacuum attached to it. DNA and beads are then kept on the comb surface, and the comb is moved through the containers 1 to 3 following the previously described order with the vacuum still on until the liquid present in each container is sucked up. Then, the comb is placed over the optical plate without touching the primer mix, and the vacuum is turned off. The comb is gently agitated to help release the DNA and the beads that are going to be mixed with the primer. The optical plate is heated to 80 °C for 2 minutes. CDT tips are filled with the nucleotides, enzyme and substrate (Qiagen) and the optical plate is loaded to the machine. The PyroMark software was used to calculate the allelic ratio for the different analyzed SNPs, in all samples, Doxycycline, No Doxycycline and Doxycycline plus PRC1 inhibitor. Those percentages were later used for the calculation of percentage of gene silencing.

### RNA FISH and Immunofluorescence (IF)

Cells with the different constructs were grown for 5 days on coverslips in Petri Dish 100×1mm X-plates, medium with Doxycycline (1ul per ml) changed daily. Next, cells were washed with PBS and fixed with the following protocol: the coverslips were put in Coplin jars with CSK buffer to rinse, then they were left for 8 minutes in CSK buffer plus Triton-X (Sigma-Aldrich). In a third step, the coverslips were moved to 4% PFA for another 8 minutes and they were rinsed in 70% ethanol (with DEPC treated H_2_O). They were finally stored in snap-cap vials containing 70% ethanol at 4 C. The RNA FISH and IF protocols were done subsequently. First, the coverslips were rinsed in 1x PBS for several seconds. They were incubated for 20 minutes facing down with 100 ul of PBT (1% (wt/vol) BSA, 0.1% (vol/vol) Tween-20 (Sigma-Aldrich) in PBS plus Ribolock (ThermoFisher), on Parafilm, an extra sheet of Parafilm was added above and sealed around each coverslip, creating some kind of individual envelope. Next, the coverslips were moved to another Parafilm envelope with 100 μl of primary antibody solution in PBT with 0.4 U μl Ribolock for a 4 to 6 hours period of incubation at room temperature, alternatively, the cells can be incubated overnight at 4 °C. The RNA probes were prepared using the Nick translation DNA labelling system 2.0 (Enzo). A mix of 6 ul of nick translated RNA probe with 12ul of Human Cot-1 (ThermoFisher) and 2ul of salmon testes (ThermoFisher) per coverslip is dried in a speedvac for at least 1.5 hours. Following the incubation with the primary antibody, the coverslips are washed with 0.1% Tween-20 in PBS for 5 min at room temperature, three times. Then, the cells were incubated with 1 ul of secondary antibody in 100 ul PBT with 0.4 U (1ul) Ribolock for 0.5–1 h at room temperature in dark. From this point, the coverslips were kept in the dark. The cells were washed again three times with PBS plus Tween-20 (Sigma-Aldrich), and then fixed with paraformaldehyde for 5 minutes and a final PBS wash for another 5 minutes at room temperature. Before initiating the FISH, the coverslips were dehydrated with ethanol at 70%, 80% and 100%, 2 minutes in each concentration. Next, they were air-dried, meanwhile the probe was resuspended in 10 ul of deionized formamide (Sigma-Aldrich) and denatured at 80 °C for 10 minutes, after that 10 ul of hybridization buffer (20% BSA and 20% Dextran Sulfate in 4x SSC) was added and gently mixed. The total volume (20 ul) was placed over Parafilm, and the coverslips were carefully put facing the cells down, it is important to not have air bubbles and again create a Parafil envelope. The coverslips were incubated overnight at 37 °C. The next day the coverslips were rinsed in a Coplin jar with deionized formamide (5 ml) combined with 4x SSC (Invitrogen) for 20 minutes at 37 °C. A second rinse was done with 2x SSC at 37 °C for 20 minutes and a final rinse with 1x SSC at room temperature for another 20 minutes. The cells were then incubated with a mix of methanol and DAPI (0.1 ug/ml) at 37 °C for 15 minutes. A final rinse with methanol was performed. The coverslips were mounted on glass slides using one drop of Vetashield and with the cells facing down. The slides are stored at −20 °C. The analysis of the cells was performed using a confocal fluorescence microscope (Leica DMI6000 B). Pictures for the blue channel (DAPI), green channel (XIST) and red channel (protein/mark) were taken and analyzed using the image processing program ImageJ (Fiji). The analysis of the cells was blinded. The images from the three channels were merged, and then a line was drawn over the XIST RNA cloud and crossed all the nucleus avoiding the nucleolar region. Selecting the BAR plugin an Excel sheet with numbers representing the pixel intensity for each channel is generated for each cell and subsequently analyzed by calculating the Z score for 50 cells per construct/clone/mark. Z score calculation lets us find the intensity in pixels for the protein/mark (in red) exactly in the place where the XIST RNA cloud is located (in green), by comparing this intensity with the rest of the signal present for both, protein and XIST, in the rest of the nucleus. A Z score of 1 means one standard deviation more for the signal intensity for the protein where XIST was seen, therefore confirming in a quantitative way the co-localization of protein/mark and XIST.

### PRC1 Inhibition

The cells were grown on coverslips for 5 days, adding Doxycycline (1 ul per ml), in a second set (per construct/per mark) in addition to Doxycycline, PRT4165 (Sigma-Aldrich) was added, different concentrations were tested (25uM, 40 uM, 45 uM and 50 uM) and the 45uM concentration was chosen. In a previous work in the laboratory, PRT4165 used in this range of concentrations had led to a reduction of ~50% of protein levels, demonstrated by Western Blot ^18^. The cells were then fixed and IF FISH for UbH2A, SMCHD1, H2K27me3 and MACROH2A was done as already described. The analysis of 30 cells by construct/mark was also blind.

### RNA Immunoprecipitation Assay

After five days of Doxycycline induction the cells were rinsed with 10 ml of cold PBS and then the PBS was removed. The cells were UV irradiated with 400mJ/cm^2^ at 254 nm using the UV Stratalinker crosslinker 1800 (STRATAGENE). The crosslinked cells were collected with a scraper and a pellet was obtained. The cells were resuspended with 200 ul of SDS Buffer (for 30 ml stock 1.5 ml of 1M Tris-HCl pH 8, 4.5 ml of 1M NaCl, 60 ul of 0.5 M EDTA, 3 ml of 10% SDS, 3ml of 10% Triton and 17.3ml of DEPC H_2_O. To prepare 500 ul of working solution, 489 ul of the previous mix was added to 0.5 ul of 1M DTT, 10 ul of Protease Inhibitor Cocktail (Roche; 1 tablet diluted in 1 ml of DEPC H_2_O), and 0.5 ul of Ribolock). The cells were incubated on ice for 10 minutes and sonicated with an S220 Focused-Ultrasonicator (Covaris), using the software SonoLab 7.2 with the following program 10 repeats of treatment with a Peak Power= 200, Cycles/Burst=50, Duty Factor=10, duration 20 seconds; alternated with a treatment of a Peak Power=2.5, Cycles/Burst= 50, Duty Factor=0.1, duration 20 seconds. Total Run time: 06 minutes 40 seconds. Then the cells received treatment with 0.5 ul of DNase (Roche) and 20 ul of 5x Buffer for 20 minutes at 37 °C. The sonication was repeated with the same program. The lysate was 10-fold diluted with the Dilution Buffer (for 30 ml stock of 1.5 ml of 1M Tris-HCl pH 8, 4.5 ml of 1M NaCl, 60 ul of 0.5 M EDTA, 3ml of 10% Triton and 20.3ml of DEPC H_2_O. To prepare 4 ml of the working solution 3.9 ml of the previous mix was added to 4 ul of 1M DTT, 80 ul of Protease Inhibitor Cocktail 50x tablets (Roche), and 4 ul of Ribolock) and spun at full speed for 30 minutes at 4 °C. Meanwhile, the antibodies were prepared by mixing 3 ug of antibody HNRNPK and IgG (as a negative control in a separate tube) with 650 ul of Dilution Buffer and 50 ul of Surebeads Protein G magnetic Beads (BioRad) and left in rotation for 30 minutes at 4 °C. Next, 300 ul of the lysate was added to each tube with the antibodies and the volume was adjusted to ~1 ml with Dilution Buffer. The tubes were left in rotation for 2 hours to overnight at 4 °C. Additionally, 1/10 of the lysate was saved and kept at −80 °C and saved as 10% input. Then, the samples with antibodies were washed, the tubes were held with a Magnetic Separator for Microcentrifuge Tubes (Sigma-Aldrich), when the liquid looked clear it was discarded and the beads were washed two times with 700 ul of High-Salt Buffer (for 30 ml stock 600 ul of 1M Tris-HCl pH 8, 15 ml of 1M NaCl, 60 ul of 0.5 M EDTA, 300 ul of 10% SDS, 3 ml of 10% Triton and 10.4 ml of DEPC H_2_O. To prepare 3 ml of the working solution 2934 ul of the previous mix was added to 3 ul of 1M DTT, 60 ul of Protease Inhibitor Cocktail 50x tablets (Roche), and 3 ul of Ribolock). Next, three subsequent washes with 700 ul of Low-Salt Buffer (for 30 ml stock 600 ul of 1M Tris-HCl pH 8, 4.5 ml of 1M NaCl, 60 ul of 0.5 M EDTA, 300 ul of 10% SDS, 3 ml of 10% Triton and 20.9 ml of DEPC H_2_O. To prepare 4.5 ml of working solution, 4401 ul of the previous solution was added to 4.5 ul of 1M DTT, 90 ul of Protease Inhibitor Cocktail 50x tablets (Roche), and 4.5 ul of Ribolock). The beads were then resuspended in 200 ul Elution Buffer (for 10 ml stock 100 ul of 1M Tris-HCl pH 8, 20 ul of 0.5 M EDTA, 500 ul of 10% SDS, and 9.4 ml of DEPC H_2_O. To prepare 500 ul of working solution, 593.4 ul of the previous mix was added to 5ul of Proteinase K 20mg/ml (Ambion), and 0.5 ul of Ribolock) and incubated for 1 hour at 37 °C. Using again the Magnetic Separator the beads were discarded and the supernatant transferred to new tubes. The DNA was precipitated with TRIzol (Invitrogen) and a second DNAse (Roche) treatment was performed.

## Acknowledgements

We would like to thank the Louis Lefebvre lab for letting us use the microscope for the IF FISH pictures. To BioRender.com for the generation of the figure 4C. The grant support for the research was from the Canadian Institutes of Health Research (PJT-156048). To the UBC Four Year Doctoral Fellowship.

## Abbreviations

(Xa): Active X
(DOX): Doxycycline
(FISH): Fluorescence in situ hybridization
(FSHD2): Facioscapulohumeral muscular dystrophy type 2
(hESC): Human embryonic stem cells
(IF): Immunofluorescence
(lncRNA): Long non-coding RNA
(SMC): Structural maintenance of chromosome
(PSC): Pluripotent stem cells
(PID): Polycomb intecation domain
(PRC): Polycomb-repressive complex
(RIP): RNA immunoprecipitation
(TAD): Topologically associated domains
(XCI): X-chromosome inactivation
(XIST): X-inactive specific transcript

## References

1. Brown, C. J. et al. A gene from the region of the human X inactivation centre is expressed exclusively from the inactive X chromosome. Nature 349, 38–44 (1991).

2. van Bemmel, J. G., Mira-Bontenbal, H. & Gribnau, J. Cis- and trans-regulation in X inactivation. Chromosoma 125, 41–50 (2016).

3. Balaton, B. P., Dixon-McDougall, T., Peeters, S. B. & Brown, C. J. The eXceptional nature of the X chromosome. Hum. Mol. Genet. 27, R242–;R249 (2018).

4. Posynick, B. J. & Brown, C. J. Escape From X-Chromosome Inactivation: An Evolutionary Perspective. Frontiers in Cell and Developmental Biology 7, 241 (2019).

5. Loda, A. & Heard, E. Xist RNA in action: Past, present, and future. PLoS Genet. 15, e1008333 (2019).

6. Boeren, J. & Gribnau, J. Xist-mediated chromatin changes that establish silencing of an entire X chromosome in mammals. Curr. Opin. Cell Biol. 70, 44–50 (2021).

7. Patrat, C., Ouimette, J.-F. & Rougeulle, C. X chromosome inactivation in human development. Development 147, (2020).

8. Peñaherrera, M. S., Ma, S., Ho Yuen, B., Brown, C. J. & Robinson, W. P. X-chromosome inactivation (XCI) patterns in placental tissues of a paternally derived bal t(X;20) case. Am. J. Med. Genet. A 118A, 29–34 (2003).

9. Okamoto, I. et al. Eutherian mammals use diverse strategies to initiate X-chromosome inactivation during development. Nature 472, 370–374 (2011).

10. Petropoulos, S. et al. Single-Cell RNA-Seq Reveals Lineage and X Chromosome Dynamics in Human Preimplantation Embryos. Cell 165, 1012–1026 (2016).

11. Okamoto, I. et al. The X chromosome dosage compensation program during the development of cynomolgus monkeys. Science 374, eabd8887 (2021).

12. Chu, C. et al. Systematic discovery of Xist RNA binding proteins. Cell 161, 404–416 (2015).

13. Moindrot, B. & Brockdorff, N. RNA binding proteins implicated in Xist-mediated chromosome silencing. Semin. Cell Dev. Biol. 56, 58–70 (2016).

14. Lu, Z. et al. Structural modularity of the XIST ribonucleoprotein complex. Nat. Commun. 11, 6163 (2020).

15. Wutz, A., Rasmussen, T. P. & Jaenisch, R. Chromosomal silencing and localization are mediated by different domains of Xist RNA. Nat. Genet. 30, 167–174 (2002).

16. Dossin, F. et al. SPEN integrates transcriptional and epigenetic control of X-inactivation. Nature 578, 455–460 (2020).

17. McHugh, C. A. et al. The Xist lncRNA interacts directly with SHARP to silence transcription through HDAC3. Nature 521, 232–236 (2015).

18. Dixon-McDougall, T. & Brown, C. J. Independent domains for recruitment of PRC1 and PRC2 by human XIST. PLoS Genet. 17, e1009123 (2021).

19. Lu, Z. et al. Structural modularity of the XIST ribonucleoprotein complex. Nat. Commun. 11, 6163 (2020).

20. Nesterova, T. B. et al. Characterization of the genomic Xist locus in rodents reveals conservation of overall gene structure and tandem repeats but rapid evolution of unique sequence. Genome Res. 11, 833–849 (2001).

21. Ridings-Figueroa, R. et al. The nuclear matrix protein CIZ1 facilitates localization of Xist RNA to the inactive X-chromosome territory. Genes Dev. 31, 876–888 (2017).

22. Wang, Z. et al. The emerging roles of hnRNPK. J. Cell. Physiol. 235, 1995–2008 (2020).

23. Nakamoto, M. Y., Lammer, N. C., Batey, R. T. & Wuttke, D. S. hnRNPK recognition of the B motif of Xist and other biological RNAs. Nucleic Acids Res. 48, 9320–9335 (2020).

24. Lange, L. et al. A de novo frameshift in HNRNPK causing a Kabuki-like syndrome with nodular heterotopia. Clin. Genet. 90, 258–262 (2016).

25. Pintacuda, G. et al. hnRNPK Recruits PCGF3/5-PRC1 to the Xist RNA B-Repeat to Establish Polycomb-Mediated Chromosomal Silencing. Mol. Cell 68, 955–969.e10 (2017).

26. Almeida, M., Bowness, J. S. & Brockdorff, N. The many faces of Polycomb regulation by RNA. Curr. Opin. Genet. Dev. 61, 53–61 (2020).

27. Sunwoo, H., Colognori, D., Froberg, J. E., Jeon, Y. & Lee, J. T. Repeat E anchors Xist RNA to the inactive X chromosomal compartment through CDKN1A-interacting protein (CIZ1). Proc. Natl. Acad. Sci. U. S. A. 114, 10654–10659 (2017).

28. Beletskii, A., Hong, Y. K., Pehrson, J., Egholm, M. & Strauss, W. M. PNA interference mapping demonstrates functional domains in the noncoding RNA Xist. Proc. Natl. Acad. Sci. U. S. A. 98, 9215–9220 (2001).

29. Hoki, Y. et al. A proximal conserved repeat in the Xist gene is essential as a genomic element for X-inactivation in mouse. Development 136, 139–146 (2009).

30. Yamada, N. et al. Xist Exon 7 Contributes to the Stable Localization of Xist RNA on the Inactive X-Chromosome. PLoS Genet. 11, e1005430 (2015).

31. Kolpa, H. J., Fackelmayer, F. O. & Lawrence, J. B. SAF-A Requirement in Anchoring XIST RNA to Chromatin Varies in Transformed and Primary Cells. Developmental cell vol. 39 9–10 (2016).

32. Sakaguchi, T. et al. Control of Chromosomal Localization of Xist by hnRNP U Family Molecules. Developmental cell vol. 39 11–12 (2016).

33. Xiao, J. et al. Mutations in CIZ1 cause adult onset primary cervical dystonia. Ann. Neurol. 71, 458–469 (2012).

34. Pandya-Jones, A., Markaki, Y., Serizay, J. & Chitiashvili, T. A protein assembly mediates Xist localization and gene silencing. Nature (2020).

35. Sofi, S. et al. Prion-like domains drive CIZ1 assembly formation at the inactive X chromosome. J. Cell Biol. 221, (2022).

36. Blewitt, M. E. et al. SmcHD1, containing a structural-maintenance-of-chromosomes hinge domain, has a critical role in X inactivation. Nat. Genet. 40, 663–669 (2008).

37. Jansz, N. et al. Smchd1 Targeting to the Inactive X Is Dependent on the Xist-HnrnpK-PRC1 Pathway. Cell Rep. 25, 1912–1923.e9 (2018).

38. Nozawa, R.-S. et al. Human inactive X chromosome is compacted through a PRC2-independent SMCHD1-HBiX1 pathway. Nat. Struct. Mol. Biol. 20, 566–573 (2013).

39. Wang, C.-Y., Jégu, T., Chu, H.-P., Oh, H. J. & Lee, J. T. SMCHD1 Merges Chromosome Compartments and Assists Formation of Super-Structures on the Inactive X. Cell 174, 406–421.e25 (2018).

40. Hamel, J. & Tawil, R. Facioscapulohumeral Muscular Dystrophy: Update on Pathogenesis and Future Treatments. Neurotherapeutics 15, 863–871 (2018).

41. Wang, C.-Y., Brand, H., Shaw, N. D., Talkowski, M. E. & Lee, J. T. Role of the Chromosome Architectural Factor SMCHD1 in X-Chromosome Inactivation, Gene Regulation, and Disease in Humans. Genetics 213, 685–703 (2019).

42. Jiang, J. et al. Translating dosage compensation to trisomy 21. Nature 500, 296–300 (2013).

43. Kelsey, A. D. et al. Impact of flanking chromosomal sequences on localization and silencing by the human non-coding RNA XIST. Genome Biol. 16, 208 (2015).

44. Dixon-McDougall, T. & Brown, C. J. Multiple distinct domains of human XIST are required to coordinate gene silencing and subsequent heterochromatin formation. Epigenetics Chromatin 15, 6 (2022).

45. Minks, J., Baldry, S. E., Yang, C., Cotton, A. M. & Brown, C. J. XIST-induced silencing of flanking genes is achieved by additive action of repeat a monomers in human somatic cells. Epigenetics Chromatin 6, 23 (2013).

46. Chapman, A. G., Cotton, A. M., Kelsey, A. D. & Brown, C. J. Differentially methylated CpG island within human XIST mediates alternative P2 transcription and YY1 binding. BMC Genet. 15, 89 (2014).

47. Nishioka, K. et al. PR-Set7 is a nucleosome-specific methyltransferase that modifies lysine 20 of histone H4 and is associated with silent chromatin. Mol. Cell 9, 1201–1213 (2002).

48. Sunwoo, H., Colognori, D., Froberg, J. E., Jeon, Y. & Lee, J. T. Repeat E anchors Xist RNA to the inactive X chromosomal compartment through CDKN1A-interacting protein (CIZ1). Proc. Natl. Acad. Sci. U. S. A. 114, 10654–10659 (2017).

49. Alchanati, I. et al. The E3 ubiquitin-ligase Bmi1/Ring1A controls the proteasomal degradation of Top2alpha cleavage complex - a potentially new drug target. PLoS One 4, e8104 (2009).

50. Dominguez, D. et al. Sequence, Structure, and Context Preferences of Human RNA Binding Proteins. Mol. Cell 70, 854–867.e9 (2018).

51. Moindrot, B. & Brockdorff, N. RNA binding proteins implicated in Xist-mediated chromosome silencing. Semin. Cell Dev. Biol. 56, 58–70 (2016).

52. Tjalsma, S. J. D. et al. H4K20me1 and H3K27me3 are concurrently loaded onto the inactive X chromosome but dispensable for inducing gene silencing. EMBO Rep. 22, e51989 (2021).

53. Wang, C.-Y., Colognori, D., Sunwoo, H., Wang, D. & Lee, J. T. PRC1 collaborates with SMCHD1 to fold the X-chromosome and spread Xist RNA between chromosome compartments. Nat. Commun. 10, 2950 (2019).

54. Colognori, D., Sunwoo, H., Kriz, A. J., Wang, C.-Y. & Lee, J. T. Xist Deletional Analysis Reveals an Interdependency between Xist RNA and Polycomb Complexes for Spreading along the Inactive X. Mol. Cell 74, 101–117.e10 (2019).

55. Bousard, A. et al. The role of Xist-mediated Polycomb recruitment in the initiation of X-chromosome inactivation. EMBO Rep. 20, e48019 (2019).

56. Cerase, A., Calabrese, J. M. & Tartaglia, G. G. Phase separation drives X-chromosome inactivation. Nat. Struct. Mol. Biol. 29, 183–185 (2022).

57. Yu, B. et al. B cell-specific XIST complex enforces X-inactivation and restrains atypical B cells. Cell 184, 1790–1803.e17 (2021).

58. Chow, J. C. et al. Inducible XIST-dependent X-chromosome inactivation in human somatic cells is reversible. Proceedings of the National Academy of Sciences 104, 10104–10109 (2007).

