## Supplementary tables and figures for "Derivation of a minimal functional XIST by combining human and mouse interaction domains"

Table S1: List of used primers for cloning, qPCR and pyrosequencing.

| Gene/ Function | Primer Forward | Primer Reverse | Primer sequencing (pyrosequencing) |
|---|---|---|---|
| qXIST 2AS(+Not1/ AgeI) clone AF | ACCGGTGCGGCCGCTAGCCTTAGGTGAGGCACCAA | | |
| qXIST10 (with Apa/Nari) Fragment E | GGCGCCGGTACCGTCCCCCAACACCCTTTAT | | |
| C7-6 (with AgeI/ XhoI) Fragment E | ACCGGTCTCGAGACAGTGAGATAGCTGCCTAG | | |
| *qUBC* | TGTCAAGGCAAAGATCCAAGATAA | TCCAGCTGTTTTCCAGCAAA | |
| *qPCDNA5* | GCTCGTTTAGTGAACCGTCAGA | GGTCCCGGTGTCTTCTATGGA | |
| *qXIST29* | CAGCGAAGACCTGGGTGAAT | CCCTCCACTTCTTTCTCCTGACT | |
| *qHyg* | TCAGAGGTTTTCACCGTCATCAC | CACCCTAACTGACACACATTCCA | |
| *AGPAT5* | ACCTTCTCAAGCCAGTGAAATAC | GGGAAATAAACAAATTCGTTCAG (biotinylated) | CAGTGAAATACAGACTTAAT |
| *CTSB* | TCGTGCACTCTGCTAATCATG | CAGTGGGTCAGAAACAACTCC (biotinylated) | TTTACAGATTGCCTCCT |
| *DLC1* (M13) | TTGATGGCTTCCAGATTTGTAA | CGCCAGGGTTTTCCCAGTCACGACCAGATCCCAAAGCCCAGTACAC | GCTTCCAGATTTGTAAGATT |
| *STC1* | TTCATTTTAGGGGTGTTGACACA | AAAAAAATCAAACCAGGCACAGT (biotinylated) | TAGGGGTGTTGACACACCA |
| *SLC25A37* | GATTTTCTTGAGGGCTCCGTAG | CGCCAGGGTTTTCCCAGTCACGACCAGA | TTGAGGGCTCCGTAG |
| *M13* | | CAGGAAACAGCTATGAC (biotinylated) | |

Table S2: List of primary antibodies used for IF.

| Primary antibody | Company | Catalogue number | Host |
|---|---|---|---|
| Anti H3K27me3 | Active Motif | 39055 | Rabbit |
| Anti UbH2A | Sigma-Aldrich | 05-678 | Mouse |
| Anti MACROH2A | Upstate | 07-219 | Rabbit |
| Anti SMCHD1 | Abcam | Ab31865 | Rabbit |
| Anti H4K20me1 | Upstate | 07-440 | Rabbit |
| Anti H3K27Ac | Active Motif | 39133 | Rabbit |
| Anti CIZ1 | Santa Cruz Biotechnology | Sc-393021 | Mouse |

Table S3: Z score averages for all constructs and testec marks. Data presented in Figure 2.

| Construct | **H3K27me3** | **UbH2A** | **MacroH2A** | **SMCHD1** | **H4K20me1** | **H3K27Ac** | **CIZ1** |
|---|---|---|---|---|---|---|---|
| AFE 1 | -0.01 | 0.04 | 0.004 | 0.13 | 0.05 | 0.12 | 3.52 |
| AFE 2 | 0.06 | 0.16 | 0.19 | 0.01 | 0.05 | 0.14 | 3.04 |
| MiniXIST 1 | 0.54 | 0.37 | 0.39 | 1.12 | 0.52 | -0.30 | 2.82 |
| MiniXIST 2 | 0.39 | 0.42 | 0.52 | 0.79 | 0.39 | -0.38 | 3.03 |
| Full XIST 1 | 1.07 | 1.05 | 0.90 | 1.54 | 0.69 | -0.66 | 3.42 |
| Full XIST 2 | 1.27 | 0.80 | 1.32 | 1.42 | 0.95 | -0.53 | 2.51 |
| IMR-90 | 1.80 | 1.15 | 1.44 | 1.93 | 1.04 | -0.93 | 1.82 |

Table S4: Z score averages for PRC1 inhibition. Data presented in Figure 3.

| Construct | **UbH2A** | **SMCHD1** | **H3K27me3** | **MACROH2A** |
|---|---|---|---|---|
| MiniXIST | 0.39 | 0.56 | 0.53 | 0.74 |
| MiniXIST Inhibitor | 0.07 | -0.16 | 0.41 | 0.71 |
| Full XIST | 1.21 | 1.08 | 0.80 | 1.23 |
| Full XIST Inhibitor | 0.73 | 0.31 | 0.59 | 0.91 |
| IMR-90 | 1.85 | 0.31 | 1.55 | 1.96 |
| IMR-90 Inhibitor | 0.91 | 1.83 | 1.12 | 1.54 |

Table S5: Percentages of silencing per construct per gene. Data shown in Figure 1D.

| Construct | *AGPAT5* | *CTSB* | *DLC1* | *STC1* | *SLC25A37* |
|---|---|---|---|---|---|
| AFE 1-1 | 2.01% | 10.58% | 0.99% | 5.02% | 2.77% |
| AFE 1-2 | 3.52% | 6.29% | -5.55% | 2.51% | -3.37% |
| AFE 1-3 | 5.03% | 8.90% | 6.05% | 7.86% | 0% |
| AFE 2-1 | 17.50% | -0.66% | 35.90% | 5.19% | 5.37% |
| AFE 2-2 | 14.04% | 21.71% | 15.67% | 7.75% | 8.48% |

| | | | | | |
|---|---|---|---|---|---|
| AFE 2-3 | 7.02% | -0.66% | 20.80% | 11.6% | 10.49% |
| MiniXIST 1-1 | 56.88% | 41.47% | 44.46% | 38.27% | 27.97% |
| MiniXIST 1-2 | 53.13% | 43.36% | 23.18% | 40.39% | 33.20% |
| MiniXIST 1-3 | 49.38% | 31.40% | 28.37% | 35.98% | 34.61% |
| MiniXIST 2-1 | 37.60% | 52.49% | 28.76% | 49.54% | 33.49% |
| MiniXIST 2-2 | 42.40% | 52.25% | 37.38% | 52.64% | 35.64% |
| MiniXIST 2-3 | 52% | 60.55% | 46.57% | 46.10% | 31.92% |
| Full XIST 1-1 | 50.36% | 50.92% | 62.57% | 76.09% | 59.91% |
| Full XIST 1-2 | 48.20% | 67.08% | 68% | 80.34% | 62.06% |
| Full XIST 1-3 | 50.36% | 78.32% | 51.71% | 78.04% | 58.54% |
| Full XIST 2-1 | 25% | 91.09% | 54.39% | 72.29% | 55.64% |
| Full XIST 2-2 | 22.92% | 64.85% | 54.07% | 69.69% | 52.45% |
| Full XIST 2-3 | 25% | 83.15% | 74.30% | 74.39% | 58.42% |

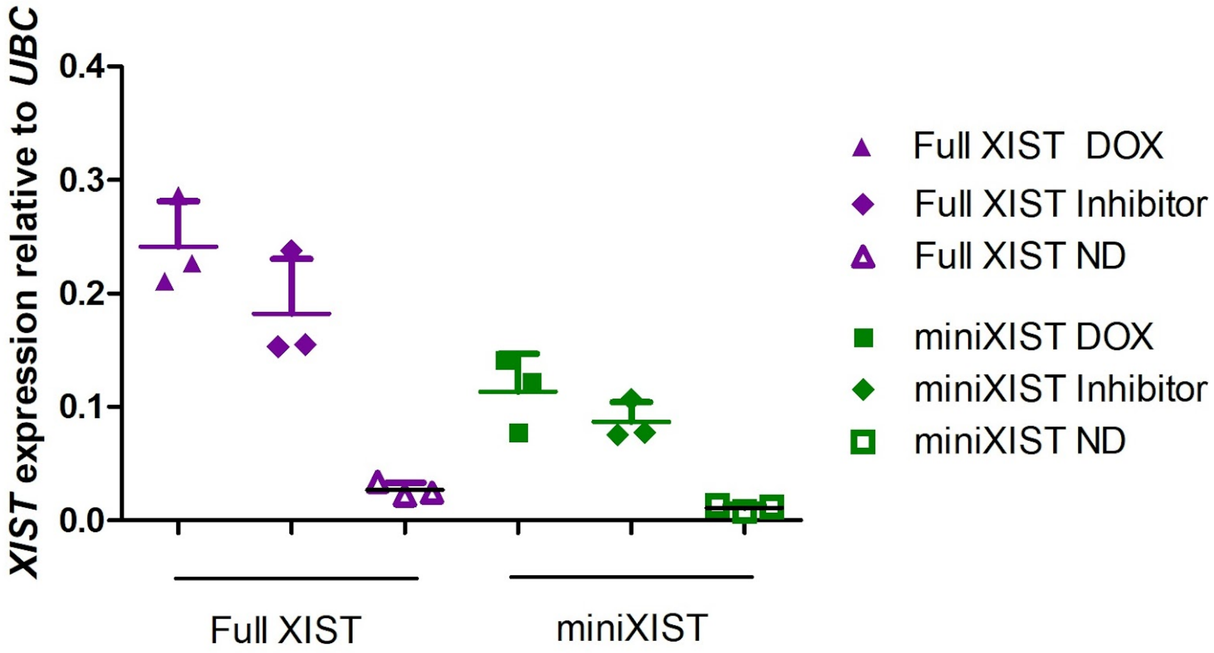


**Supplementary figure 1**: Expression of the transgenes Full XIST and miniXIST compared to *UBC* when PRT4165 is added to the cells along with Doxycycline versus cells with only Doxycycline and cells without Doxycycline.

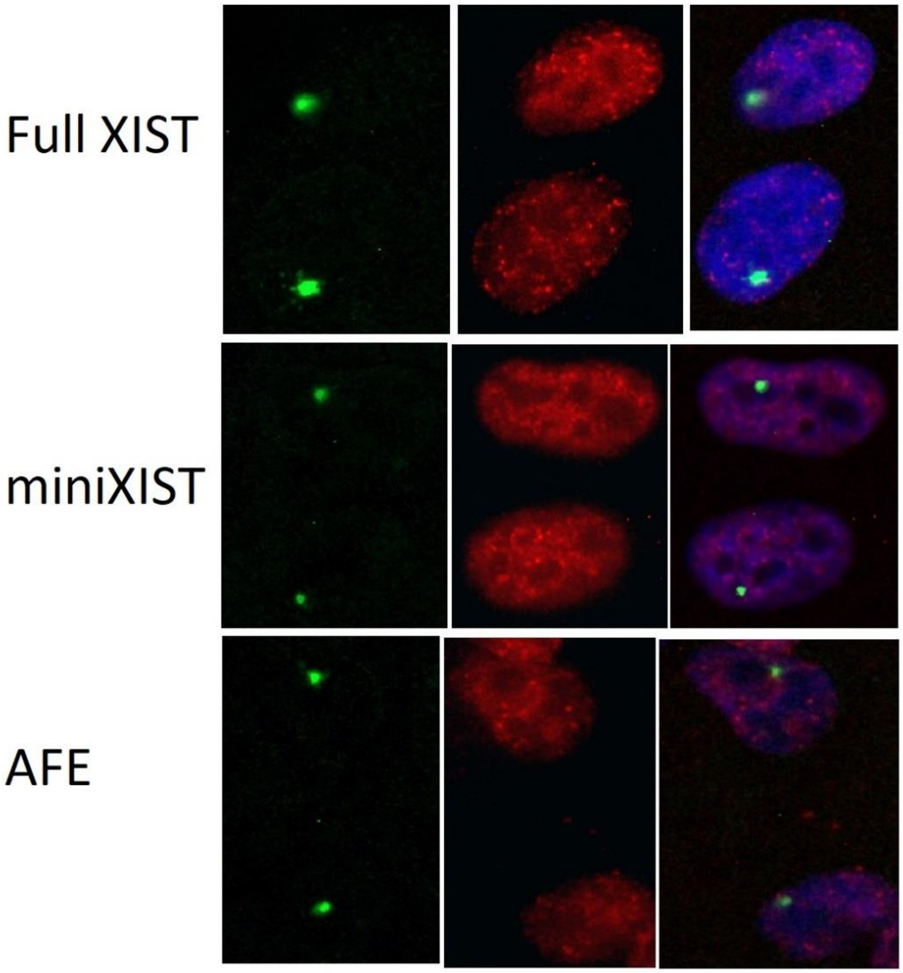


**Supplementary figure 2**: Example of IF FISH for H3K27me3 for constructs Full XIST, miniXIST and AFE.

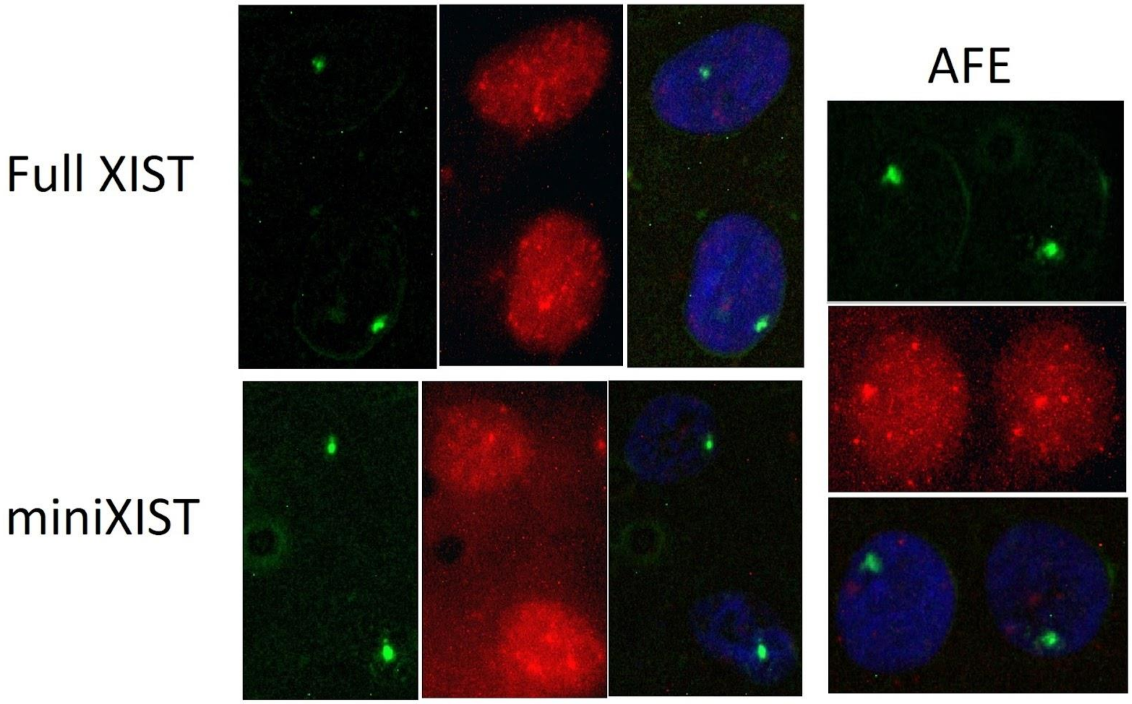


**Supplementary figure 3**: Example of IF FISH for UbH2A for constructs Full XIST, miniXIST and AFE.

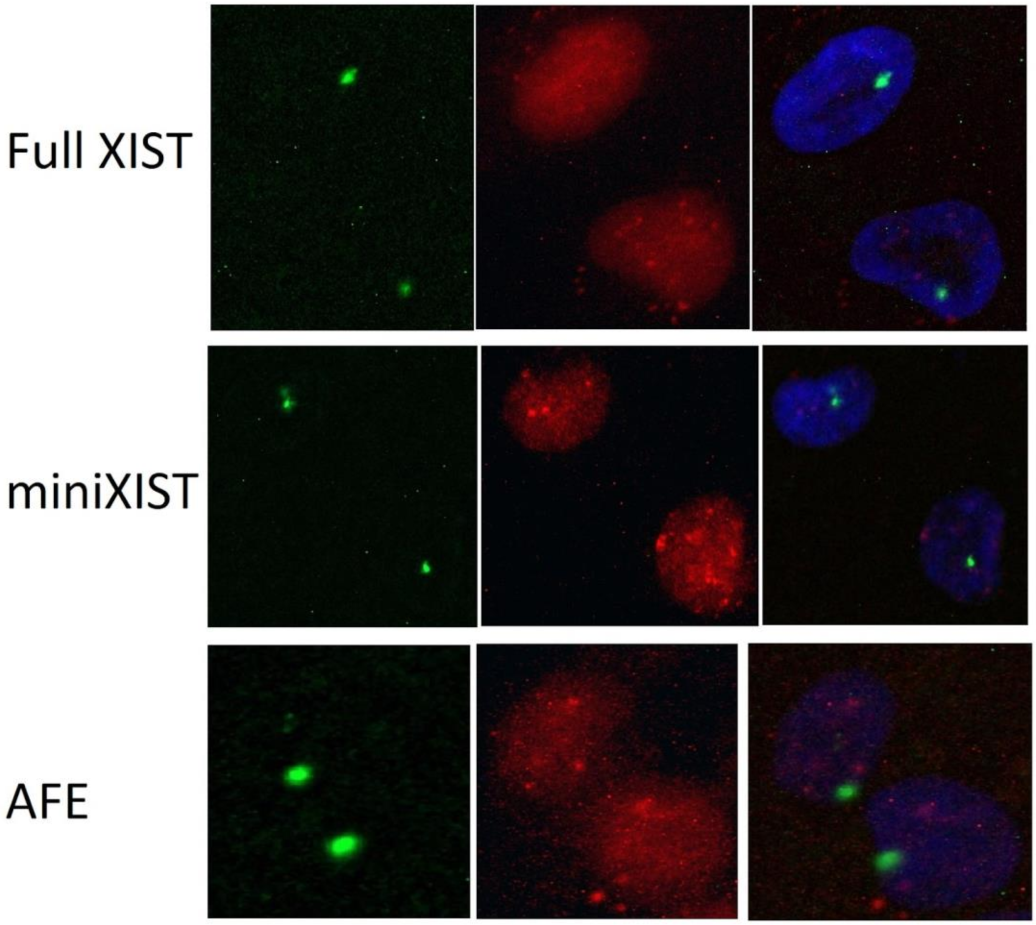


**Supplementary figure 4**: Example of IF FISH for SMCHD1 for constructs Full XIST, miniXIST and AFE.

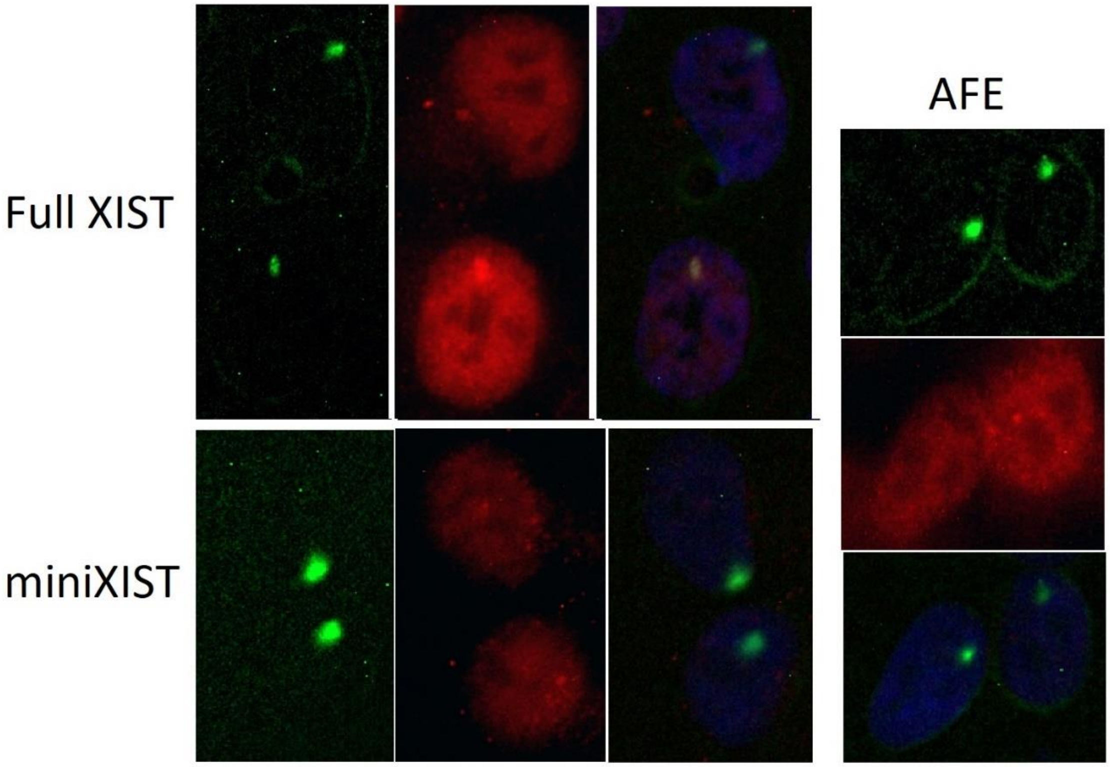


**Supplementary figure 5**: Example of IF FISH for MACROH2A for constructs Full XIST, miniXIST and AFE.

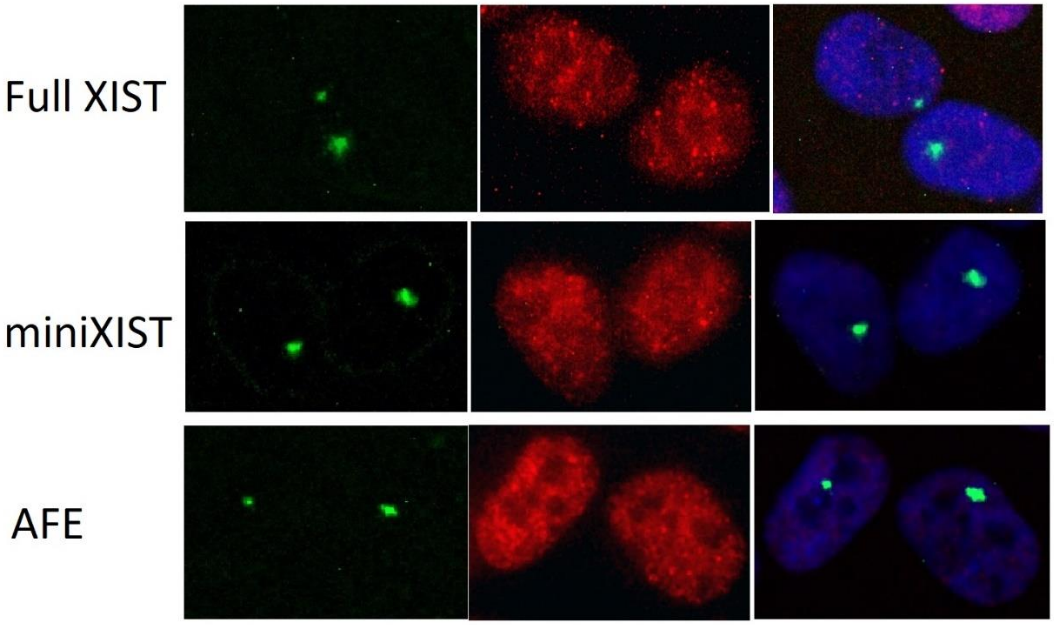


**Supplementary figure 6**: Example of IF FISH for H3K27Ac for constructs Full XIST, miniXIST and AFE.

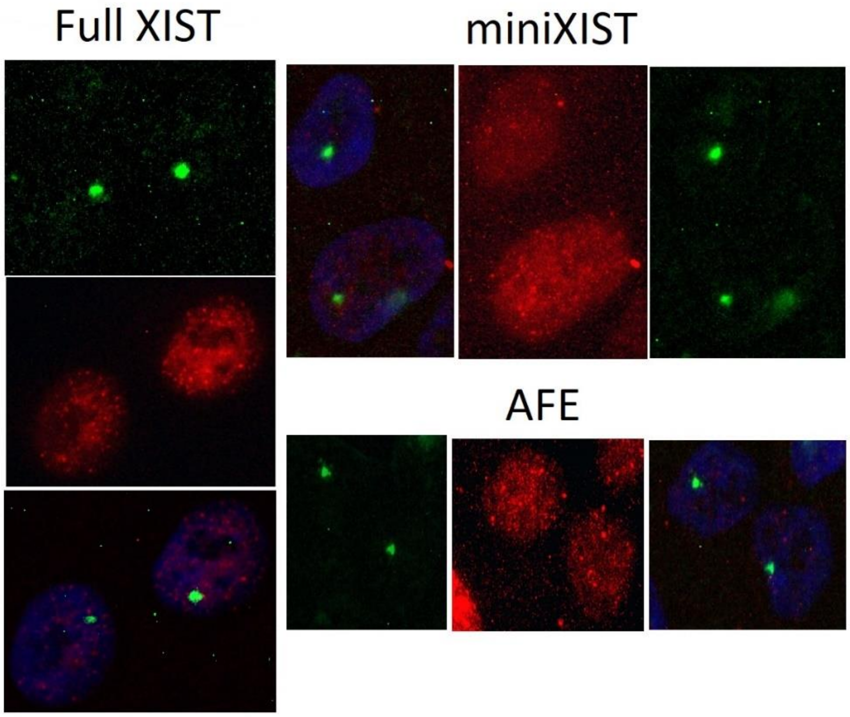


**Supplementary figure 7**: Example of IF FISH for H4K20me1 for constructs Full XIST, miniXIST and AFE.

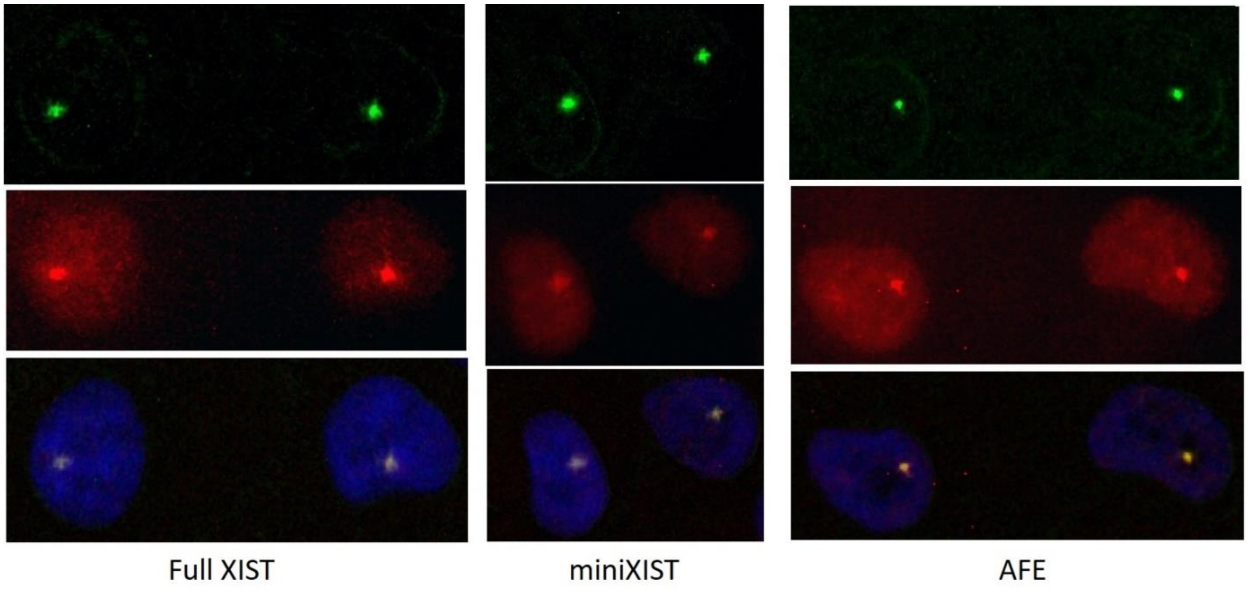


**Supplementary figure 8**: Example of IF FISH for CIZ1 for constructs Full XIST, miniXIST and AFE.
